# Mapping single-cell atlases throughout Metazoa unravels cell type evolution

**DOI:** 10.1101/2020.09.28.317784

**Authors:** Alexander J. Tarashansky, Jacob M. Musser, Margarita Khariton, Pengyang Li, Detlev Arendt, Stephen R. Quake, Bo Wang

## Abstract

Comparing single-cell transcriptomic atlases from diverse organisms can elucidate the origins of cellular diversity and assist the annotation of new cell atlases. Yet, comparison between distant relatives is hindered by complex gene histories and diversifications in expression programs. Previously, we introduced the self-assembling manifold (SAM) algorithm to robustly reconstruct manifolds from single-cell data (Tarashansky et al., 2019). Here, we build on SAM to map cell atlas manifolds across species. This new method, SAMap, identifies homologous cell types with shared expression programs across distant species within phyla, even in complex examples where homologous tissues emerge from distinct germ layers. SAMap also finds many genes with more similar expression to their paralogs than their orthologs, suggesting paralog substitution may be more common in evolution than previously appreciated. Lastly, comparing species across animal phyla, spanning mouse to sponge, reveals ancient contractile and stem cell families, which may have arisen early in animal evolution.

## Introduction

There is much ongoing success in producing single-cell transcriptomic atlases to investigate the cell type diversity within individual organisms (Regev et al., 2017). With the growing diversity of cell atlases across the tree of life (Briggs et al., 2018; Cao et al., 2019; Fincher et al., 2018; Hu et al., 2020; Musser et al., 2019; Plass et al., 2018; Siebert et al., 2019; Wagner et al., 2018), a new frontier is emerging: the use of cross-species cell type comparisons to unravel the origins of cellular diversity and uncover species-specific cellular innovations (Arendt et al., 2019; Shafer, 2019). Further, these comparisons promise to accelerate cell type annotation and discovery by transferring knowledge from well-studied model organisms to under-characterized animals.

However, recent comparative single-cell analyses are mostly limited to species within the same phylum (Baron et al., 2016; Geirsdottir et al., 2019; Sebé-Pedrós et al., 2018; Tosches et al., 2018). Comparisons across longer evolutionary distances and across phyla are challenging for two major reasons. First, gene regulatory programs diversify during evolution, diminishing the similarities in cell type specific gene expression patterns. Second, complex gene evolutionary history causes distantly related organisms to share few one-to-one gene orthologs (Nehrt et al., 2011), which are often relied upon for comparative studies (Briggs et al., 2018; Shafer, 2019). This effect is compounded by the growing evidence suggesting that paralogs may be more functionally similar than orthologs across species, due to differential gain (neo-functionalization), loss (non-functionalization), or partitioning (sub-functionalization) events among paralogs (Nehrt et al., 2011; Prince & Pickett, 2002; Stamboulian et al., 2020; Studer & Robinson-Rechavi, 2009).

Here, we present the Self-Assembling Manifold mapping (SAMap) algorithm to enable mapping single-cell transcriptomes between phylogenetically remote species. SAMap relaxes the constraints imposed by sequence orthology, using expression similarity between mapped cells to infer the relative contributions of homologous genes, which in turn refines the cell type mapping. In addition, SAMap uses a graph-based data integration technique to identify reciprocally connected cell types across species with greater robustness than previous single-cell data integration methods (Haghverdi et al., 2018; Hie et al., 2019; Polański et al., 2019; Stuart et al., 2019).

Using SAMap, we compared seven whole-body cell atlases from species spanning animal phylogeny, which have divergent transcriptomes and complex molecular homologies (**Figure 1A-B** and **Supplementary Table 1**). We began with well-characterized cell types in developing frog and fish embryos. We found broad concordance between transcriptomic signatures and ontogenetic relationships, which validated our mapping results, yet also detected striking examples of homologous cell types emerging from different germ layers. We next extended the comparison to animals from the same phylum but with highly divergent body plans, using a planarian flatworm and a parasitic blood fluke, and found one-to-one homologies even between cell subtypes. Comparing all seven species from sponge to mouse, we identified densely interconnected cell type families broadly shared across animals, including contractile and stem cells, along with their respective gene expression programs. Lastly, we noticed that homologous cell types often exhibit differential expression of orthologs and similar expression of paralogs, suggesting that the substitution and swapping of paralogs in cell types may be more common in evolution than previously appreciated. Overall, our study represents an important step towards analyzing the evolutionary origins of specialized cell types and their associated gene expression programs in animals.

**Figure 1:**
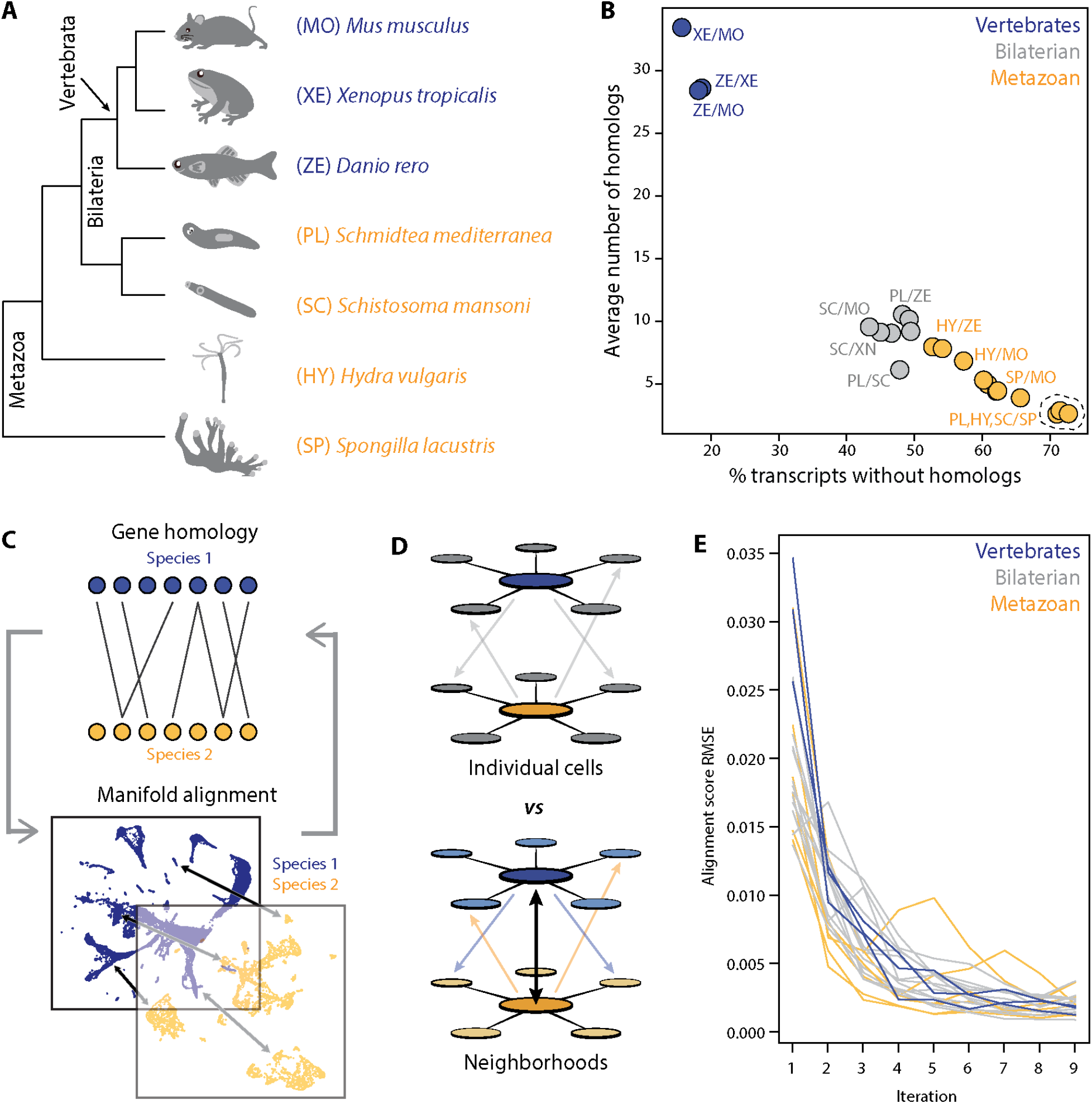
SAMap addresses challenges in mapping cell atlases of distantly related species. (A) Schematic showing the phylogenetic relationships among 7 species analyzed. (B) Challenges in mapping single-cell transcriptomes. Gene duplications cause large numbers of homologs per gene, determined by reciprocal BLAST (cut-off: e-value < 10^−6^), and frequent gene losses and the acquisition of new genes results in large fractions of transcriptomes lacking homology, which limits the amount of information comparable across species. (C) SAMap workflow. Homologous gene pairs initially weighted by protein sequence similarity are used to align the manifolds, low dimensional representations of the cell atlases. Gene-gene correlations calculated from the aligned manifolds are used to update the edge weights in the bipartite graph, which are then used to improve manifold alignment. (D) Mutual nearest neighborhoods improve the detection of cross-species mutual nearest neighbors by connecting cells that target one other’s within-species neighborhoods. (E) Convergence of SAMap is evaluated by the root mean square error (RMSE) of the alignment scores between mapped clusters in adjacent iterations for all 21 pairwise comparisons of the 7 species.

## Results

### The SAMap algorithm

SAMap iterates between two modules. The first module constructs a gene-gene bipartite graph with cross-species edges connecting homologous gene pairs, initially weighted by protein sequence similarity (**Figure 1C**). In the second module, SAMap uses the gene-gene graph to project the two single-cell transcriptomic datasets into a joint, lower-dimensional manifold representation, from which each cell’s mutual cross-species neighbors are linked to stitch the cell atlases together (**Figure 1D**). Then, using the joint manifold, the expression correlations between homologous genes are computed and used to reweight the edges in the gene-gene homology graph in order to relax SAMap’s initial dependence on sequence similarity. The new homology graph is used as input to the subsequent iteration of SAMap, and the algorithm continues until convergence, defined as when the cross-species mapping does not significantly change between iterations (**Figure 1E**).

This algorithm overcomes several challenges inherent to mapping single-cell transcriptomes between distantly related species. First, complex gene evolutionary history often results in many-to-many homologies with convoluted functional relationships (Briggs et al., 2018; Nehrt et al., 2011). SAMap accounts for this by using the full homology graph to project each dataset into both its own and its partner’s respective principal component (PC) spaces, constructed by the SAM algorithm, which we previously developed to robustly and sensitively identify cell types (Tarashansky et al., 2019). The resulting within- and cross-species projections are concatenated to form the joint space. For the cross-species projections, we translate each species’ features into those of its partner, with the expression for individual genes imputed as the weighted average of their homologs specified in the gene-gene bipartite graph. Iteratively refining the homology graph to only include positively correlated gene pairs prunes the many-to-many homologies to only include genes that are expressed in the same mapped cell types.

Second, frequent gene losses and the acquisitions of new genes result in many cell type gene expression signatures being species-specific, limiting the amount of information that is comparable across species. Restricting the analysis of each dataset to only include genes that are shared across species would result in a decreased ability to resolve cell types and subtypes with many species-specific gene signatures. SAMap solves this problem by constructing the joint space through the concatenation of within- and cross-species projections, thus encoding all genes from both species.

Lastly, the evolution of expression programs gradually diminishes the similarity between homologous cell types. To account for this effect, SAMap links cell types across species while tolerating their differences. Cells are mapped by calculating each of their *k* mutual nearest cross-species neighbors in the combined projection. To establish more robust mutual connectivity, we integrate information from each cell’s local, within-species neighborhood (**Figure 1D**), overcoming the inherent stochasticity of cross-species correlations. Two cells are thus defined as mutual nearest cross-species neighbors when their respective neighborhoods have mutual connectivity. It is important to note that the magnitude of connections is not directly calculated from their expression similarity, allowing cell types with diverged expression profiles to be tightly linked if they are among each other’s closest cross-species neighbors.

### Paralog substitutions are prevalent between homologous cell types in frog and fish

We first applied SAMap to the *Xenopus* and zebrafish atlases, which both encompass embryogenesis until early organogenesis (Briggs et al., 2018; Wagner et al., 2018). Previous analysis had linked cell types between these two organisms by matching ontogeny, thereby providing a reference for comparison. SAMap produced a combined manifold with a high degree of cross-species alignment while maintaining high resolution for distinguishing cell types in each species (**Figure 2A**). We measured the mapping strength between cell types by calculating an alignment score (edge width in **Figure 2B** and color map in **Figure 2C**), defined as the average number of mutual nearest cross-species neighbors of each cell relative to the maximum possible number of neighbors.

**Figure 2:**
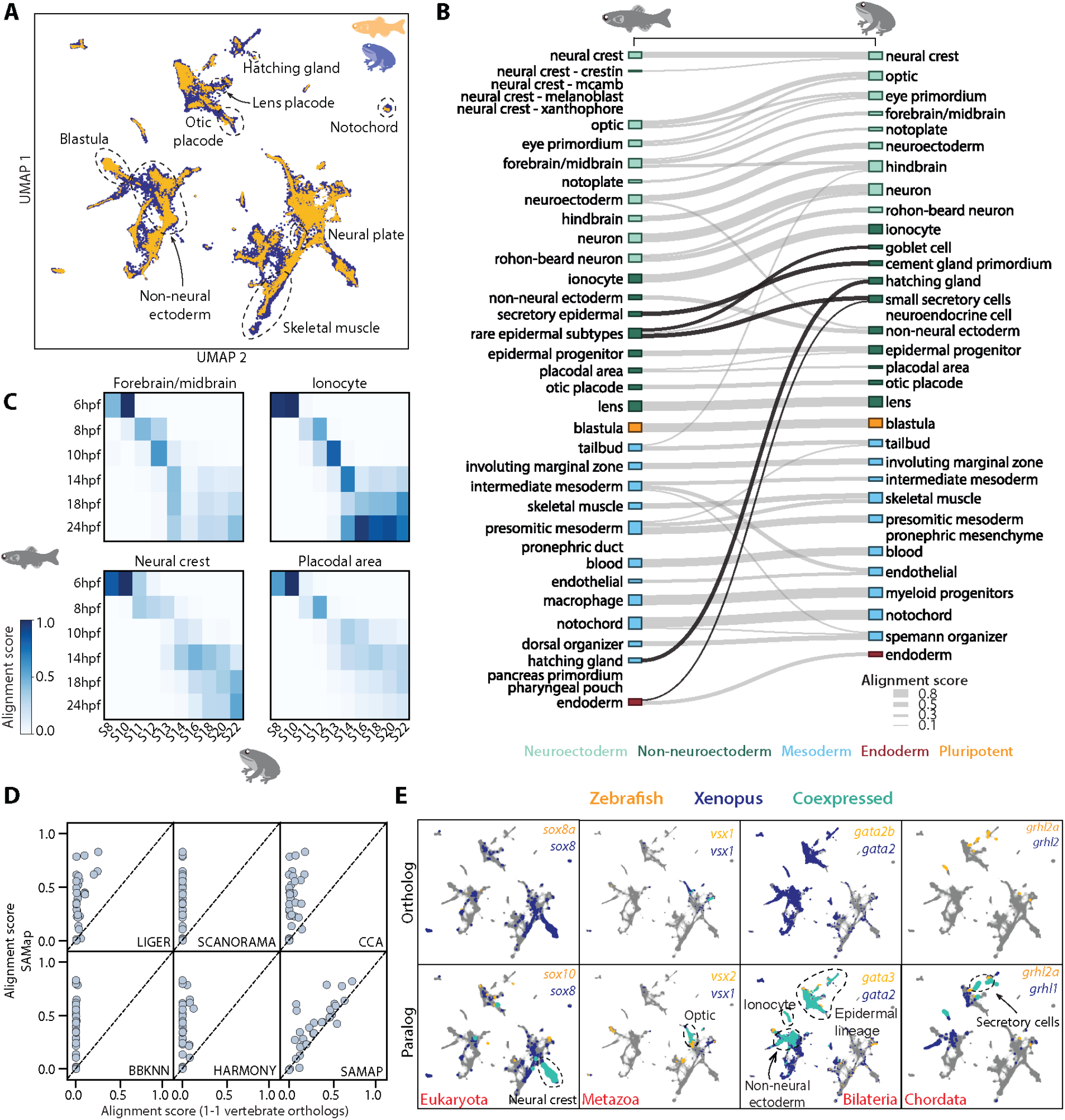
SAMap successfully maps *D. rerio* and *X. tropicalis* atlases and reveals prevalent paralog substitutions. (A) UMAP projection of the combined zebrafish (yellow) and *Xenopus* (blue) manifolds, with example cell types circled. (B) Sankey plot summarizing the cell type mappings. Edges with alignment score < 0.1 are omitted. Edges that connect developmentally distinct secretory cell types are highlighted in black. (C) Heatmaps of alignment scores between developmental time points for ionocyte, forebrain/midbrain, placodal, and neural crest lineages. X-axis: zebrafish. Y-axis: *Xenopus*. (D) SAMap alignment scores compared to those of benchmarking methods using one-to-one vertebrate orthologs as input. Each dot represents a cell type pair supported by ontogeny annotations. (E) Expression of orthologous (left) and paralogous (right) gene pairs overlaid on the combined UMAP projection. Expressing cells are color-coded by species, with those that are connected across species colored cyan. Cells with no expression are shown in gray. More examples are provided in **Figure 2 – figure supplement 3**.

SAMap revealed broad agreement between transcriptomic similarity and developmental ontogeny, linking 26 out of 27 expected pairs based on previous annotations (**Figure 2B** and **Supplementary Table 2**) (Briggs et al., 2018). The only exception is the embryonic kidney (pronephric duct/mesenchyme), potentially indicating that their gene expression programs have significantly diverged. In addition, SAMap succeeded in drawing parallels between the development of homologous cell types and matched time points along several cell lineages (**Figure 2C**). While the concordance was consistent across cell types, we noticed that the exact progression of developmental timing can vary, suggesting that SAMap can quantify heterochrony with cell type resolution.

SAMap also weakly linked several closely related cell types with different ontogeny. For example, optic cells from both species are also connected to eye primordium, frog skeletal muscles to fish presomitic mesoderm, and frog hindbrain to fish forebrain/midbrain. Notable exceptions also included mapped secretory cell types that differ in their developmental origin and even arise from different germ layers (black edges in **Figure 2B**). They are linked through a large set of genes including conserved transcription factors (e.g., *foxa1* (Dubaissi et al., 2014), *grhl* (Miles et al., 2017)) and proteins involved in vesicular protein trafficking (**Figure 2 – figure supplement 1**). This observation supports the notion that cell types may be transcriptionally and evolutionarily related despite having different developmental origins (Arendt et al., 2016).

To benchmark SAMap performance, we used Eggnog (Huerta-Cepas et al., 2019) to define one-to-one vertebrate orthologs between fish and frog and fed these gene pairs as input to several broadly used single-cell data integration methods, Seurat (Stuart et al., 2019), Liger (Welch et al., 2019), Harmony (Korsunsky et al., 2019), Scanorama (Hie et al., 2019), and BBKNN (Polański et al., 2019). We found that they failed to map the two atlases, yielding minimal alignment between them (**Figure 2D** and **Figure 2 – figure supplement 2**). We also compared the results when restricting SAMap to using the one-to-one orthologs instead of the full homology graph. Even when removing the many-to-many gene homologies and the iterative refinement of the homology graph, we identified similar, albeit weaker, cell type mappings. This suggests that, at least for the frog and fish comparison, SAMap’s performance is owed in large part to its robust, atlas stitching approach.

The key benefit of using the full homology graph is to enable the systematic identification of gene paralogs that exhibit greater similarity in expression across species than their corresponding orthologs. These events are expected to arise as the result of gene duplications followed by diversification of the resulting in-paralogs (Studer & Robinson-Rechavi, 2009). In addition, genetic compensation by transcriptional adaptation, where loss-of-function mutations are balanced by upregulation of related genes with similar sequences (El-Brolosy et al., 2019), could also result in this signature. In total, SAMap selected 8,286 vertebrate orthologs and 7,093 eukaryotic paralogs, as enumerated by Eggnog, for manifold alignment. Among these, 565 genes have markedly higher expression correlations (correlation difference > 0.3) with their paralogs than their orthologs (**Figure 2E** and **Figure 2 – figure supplement 3**), and 209 genes have orthologs that are either completely absent or lowly-expressed with no cell-type specificity (**Supplementary Table 3**), suggesting that they may have lost their functional roles at some point and were compensated for by their paralogs. We term these events as “paralog substitutions”. SAMap linked an additional 297 homologous pairs not previously annotated by orthology or paralogy, but which exhibited sequence similarity and high expression correlations (>0.5 Pearson correlation). These likely represent unannotated orthologs/paralogs or isofunctional, distantly related homologs (Gabaldón & Koonin, 2013). These results illustrate the potential of SAMap in leveraging single-cell gene expression data for pruning the networks of homologous genes to identify evolutionary substitution of paralogs and, more generally, identify non-orthologous gene pairs that may perform similar functions in the cell types within which they are expressed.

### Homologous cell types between two flatworm species with divergent body plans

To test if we can identify homologous cell types in animals with radically different body plans, we mapped the cell atlases of two flatworms, the planarian *Schmidtea mediterranea* (Fincher et al., 2018), and the trematode *Schistosoma mansoni*, which we collected recently (Li et al., 2020). They represent two distant lineages within the same phylum but have remarkably distinct body plans and autecology (Laumer et al., 2015; Littlewood & Waeschenbach, 2015). While planarians live in freshwater and are known for their ability to regenerate (Reddien, 2018), schistosomes live as parasites in humans. The degree to which cell types are conserved between them is unresolved, given the vast phenotypic differences caused by the transition from free-living to parasitic habits (Laumer et al., 2015).

SAMap revealed broad cell type homology between schistosomes and planarians. The schistosome had cells mapped to the planarian stem cells, called neoblasts, as well as most of the differentiated tissues: neural, muscle, intestine, epidermis, parenchymal, protonephridia, and *cathepsin*^+^ cells, the latter of which consists of cryptic cell types that, until now, have only been found in planarians (Fincher et al., 2018) (**Figure 3A**). These mappings are supported by both known cell type specific marker genes and numerous homologous transcriptional regulators (**Figure 3B** and **Figure 3 – figure supplement 1**).

We next determined if cell type homologies exist at the subtype level. For this, we compared the neoblasts, as planarian neoblasts are known to comprise populations of pluripotent cells and tissue-specific progenitors (Fincher et al., 2018; Zeng et al., 2018). By mapping the schistosome neoblasts to a planarian neoblast atlas (Zeng et al., 2018), we found that the schistosome has a population of neoblasts (ε-cells (Wang et al., 2018)) that cluster with the planarian’s pluripotent neoblasts, both expressing a common set of TFs (e.g., *soxp2, unc4, pax6a, gcm1*) (**Figure 3C-D**). The ε-cells are closely associated with juvenile development and lost in adult schistosomes (Wang et al., 2018), indicating pluripotent stem cells may be a transient population restricted to their early developmental stages. This is consistent with the fact that, whereas schistosomes can heal wounds, they have limited regenerative ability (Wendt & Collins, 2016). SAMap also linked other schistosome neoblast populations with planarian progenitors, including two populations of schistosome neoblasts (denoted as μ (Tarashansky et al., 2019) and μ’) to planarian muscle progenitors, all of which express *myoD*, a canonical master regulator of myogenesis (Scimone et al., 2017). These likely represent early and late muscle progenitors, respectively, as μ-cells do not yet express differentiated muscle markers such as *troponin*, whereas μ’-cells do (**Figure 3 – figure supplement 2)**.

**Figure 3:**
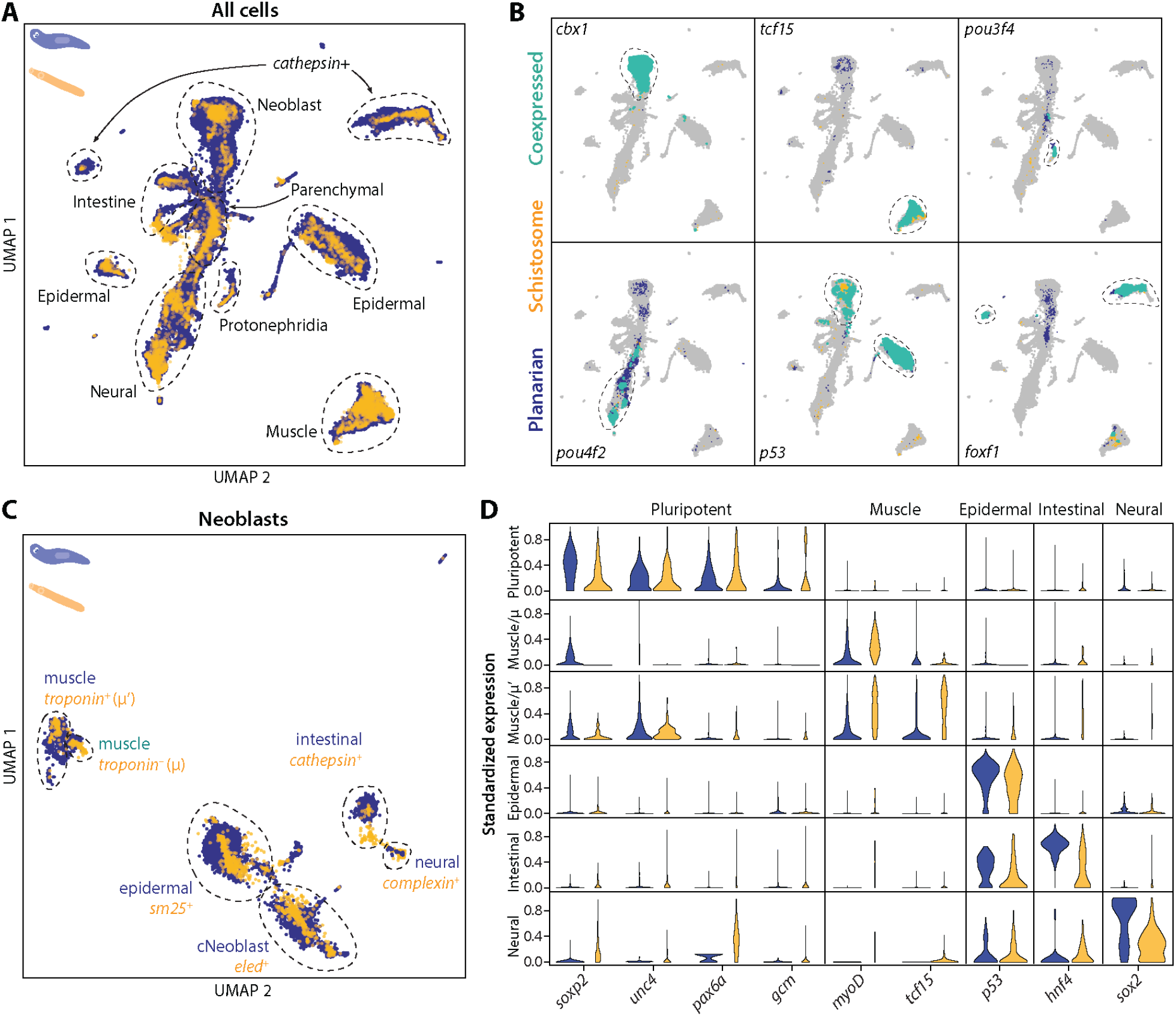
SAMap transfers cell type information from a well-annotated organism (planarian *S. mediterranea*) to its less-studied cousin (schistosome *S. mansoni*) and identifies parallel stem cell compartments. (A) UMAP projection of the combined manifolds. Tissue type annotations are adopted from the *S. mediterranea* atlas (Fincher et al., 2018). The schistosome atlas was collected from juvenile worms, which we found to contain neoblasts with an abundance comparable to that of planarian neoblasts (Li et al., 2020). (B) Overlapping expressions of selected tissue-specific TFs with expressing cell types circled. (C) UMAP projection of the aligned manifolds showing planarian and schistosome neoblasts, with homologous subpopulations circled. Planarian neoblast data is from (Zeng et al., 2018), and cNeoblasts correspond to the Nb2 population, which are pluripotent cells that can rescue neoblast-depleted planarians in transplantation experiments. (D) Distributions of conserved TF expressions in each neoblast subpopulation. Expression values are *k*-nearest-neighbor averaged and standardized, with negative values set to zero. Blue: planarian; yellow: schistosome.

### Cell type families spanning the animal tree of life

To compare cell types across broader taxonomic scales, we extended our analysis to include juvenile freshwater sponge (*Spongilla lacustris*) (Musser et al., 2019), adult *Hydra* (*Hydra vulgaris*) (Siebert et al., 2019), and mouse (*Mus musculus*) embryogenesis (Pijuan-Sala et al., 2019) atlases. In total, SAMap linked 1,051 cross-species pairs of cell types, defined by the annotations used in each respective study. 95% of the cell type pairs are supported by at least 40 enriched gene pairs, and 87% are supported by more than 100 gene pairs, indicating that SAMap does not spuriously connect cell types with limited overlap in expression profiles (**Figure 4A**).

**Figure 4:**
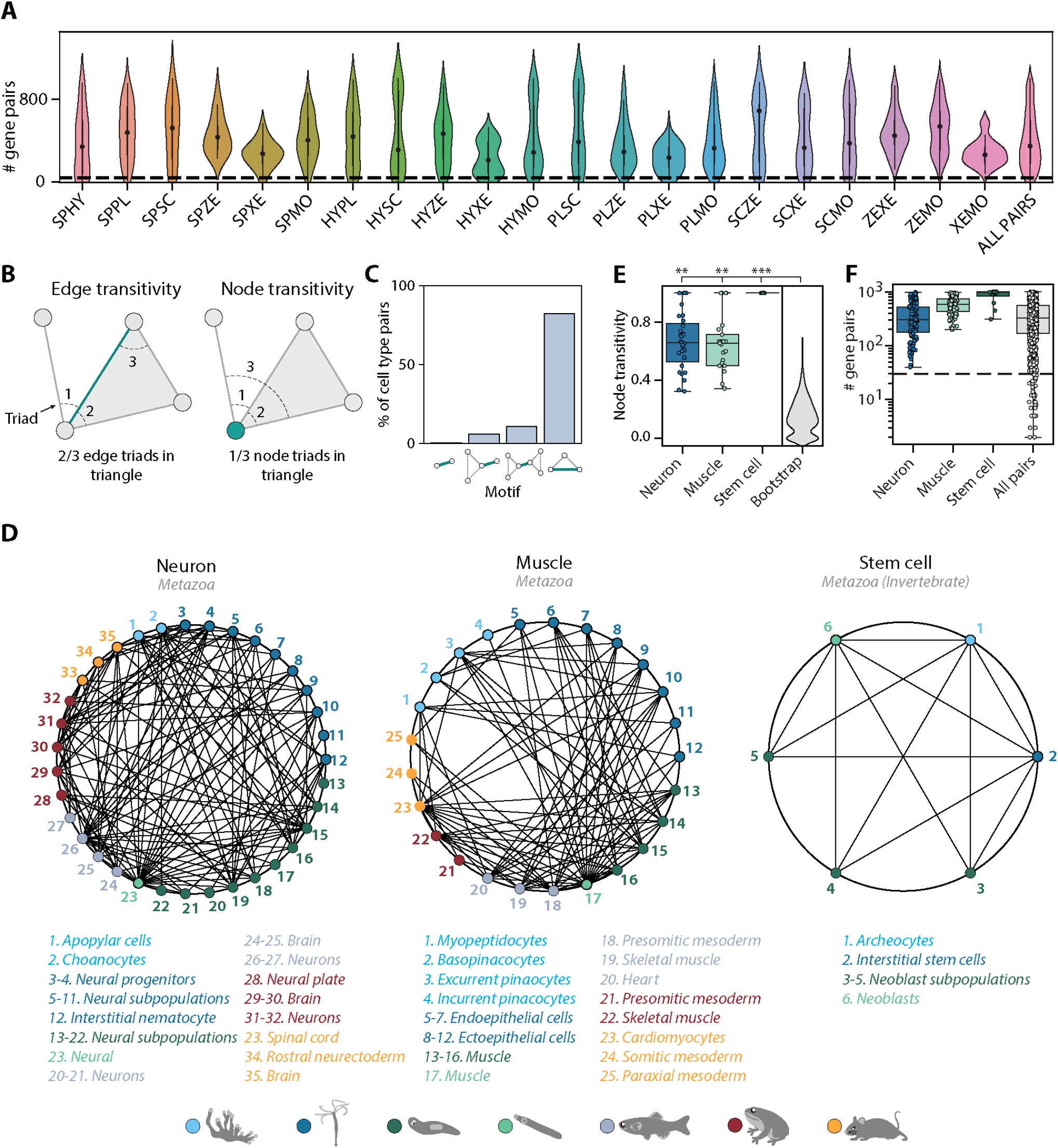
Mapping evolutionarily distant species identifies densely connected cell type groups. (A) Violin plots showing the number of enriched gene pairs in cell type mappings from all 21 pairwise mappings between the 7 species. 87% of cell type mappings have greater than 40 enriched gene pairs (dotted line). Species acronyms are the same as in **Figure 1A**. (B) Schematic illustrating edge (left) and node (right) transitivities, defined as the fraction of triads (set of three connected nodes) in closed triangles. (C) The percentage of cell type pairs that are topologically equivalent to the green edge in each illustrated motif. (D) Network graphs showing highly connected cell type families. Each node represents a cell type, color-coded by species (detailed annotations are provided in **Supplementary Table 5**). Mapped cell types are connected with an edge. (E) Boxplot showing the median and interquartile ranges of node transitivities for highly connected cell type groups. For all box plots, the whiskers denote the maximum and minimum observations. The average node transitivity per group is compared to a bootstrapped null transitivity distribution, generated by repeatedly sampling subsets of nodes in the cell type graph and calculating their transitivities. *p < 5×10^−3^, ** p < 5×10^−5^, ***p < 5×10^−7^. (F) Boxplot showing the median and interquartile ranges of the number of enriched gene pairs in highly connected cell type groups. All cell type connections in these groups have at least 40 enriched gene pairs (dotted line).

We next extended the notion of cell type pairs to cell type trios, as mapped cell types gain additional support if they share transitive relationships to other cell types through independent mappings, forming cell type triangles among species. The transitivity of a cell type pair (edge) or a cell type (node) can be quantified as the fraction of triads to which they belong that form triangles (**Figure 4B**). The majority (81%) of cell type pairs have non-zero transitivity independent of alignment score and the number of enriched gene pairs (**Figure 4 – figure supplement 1-2**). Cell type pairs with fewer than 40 enriched gene pairs tend to have lower (<0.4) transitivity (**Figure 4 – figure supplement 2**). In addition, 16% of mapped cell type pairs have zero edge transitivity but non-zero node transitivity, meaning that at least one of the cell types connects to only a single member of an interconnected cell type group (**Figure 4C**). Such edges may be of lower confidence as they should connect to other members of the same group and are thus excluded from downstream analysis.

Among the interconnected groups of cell types, we identified families of contractile cells and neural cells (**Figure 4D**). Both cell type families are highly transitive compared to the overall graph transitivity (bootstrap p-value < 1×10^−5^), meaning that their constituent cell types have more transitive edges within the group than outside the group (**Figure 4E**). In addition, the dense, many-to-many connections within the contractile and neural families are each supported by at least 40 enriched gene pairs (**Figure 4F**). Consistent with the nerve net hypothesis suggesting a unified origin of neural cell types (Tosches & Arendt, 2013), the neural family includes vertebrate brain tissues, both bilaterian and cnidarian neurons, cnidarian nematocytes that share the excitatory characteristics of neurons (Weir et al., 2020), and *Spongilla* choanocytes and apopylar cells, both of which are not considered as neurons but have been shown to express postsynaptic-like scaffolding machinery (Musser et al., 2019; Wong et al., 2019). The contractile family includes myocytes in bilaterian animals, *Hydra* myoepithelial cells that are known to have contractile myofibrils (Buzgariu et al., 2015), and sponge pinacocytes and myopeptidocytes, both of which have been implicated to play roles in contractility (Musser et al., 2019; Sebé-Pedrós et al., 2018). In contrast to the families encompassing all seven species, we also found a fully interconnected group that contains invertebrate pluripotent stem cells, including planarian and schistosome neoblasts, *Hydra* interstitial cells, and sponge archeocytes (Alié et al., 2015). The lack of one-to-one connections across phyla is in keeping with recent hypotheses that ancestral cell types diversified into families of cell types after speciation events (Arendt et al., 2016, 2019). Our findings thus suggest that these cell type families diversified early in animal evolution.

### Transcriptomic signatures of cell type families

The high interconnectedness between cell types across broad taxonomic scales suggests that they should share ancestral transcriptional programs (Arendt et al., 2016; Tosches et al., 2018). SAMap identified broad transcriptomic similarity between bilaterian and non-bilaterian contractile cells that extends beyond the core contractile apparatus. It links a total of 23,601 gene pairs, connecting 5,471 unique genes, which are enriched in at least one contractile cell type pair. Performing functional enrichment analysis on these genes, we found cytoskeleton and signal transduction functions to be enriched (p-value < 10^−3^) based on the KOG functional classifications (Tatusov et al., 2003) assigned by Eggnog (**Figure 5A**). These genes include orthology groups spanning diverse functional roles in contractile cells, including actin regulation, cell adhesion and stability, and signaling (**Figure 5B** and **Supplementary Table 4**), indicating that contractile cells were likely multifunctional near the beginning of animal evolution.

**Figure 5:**
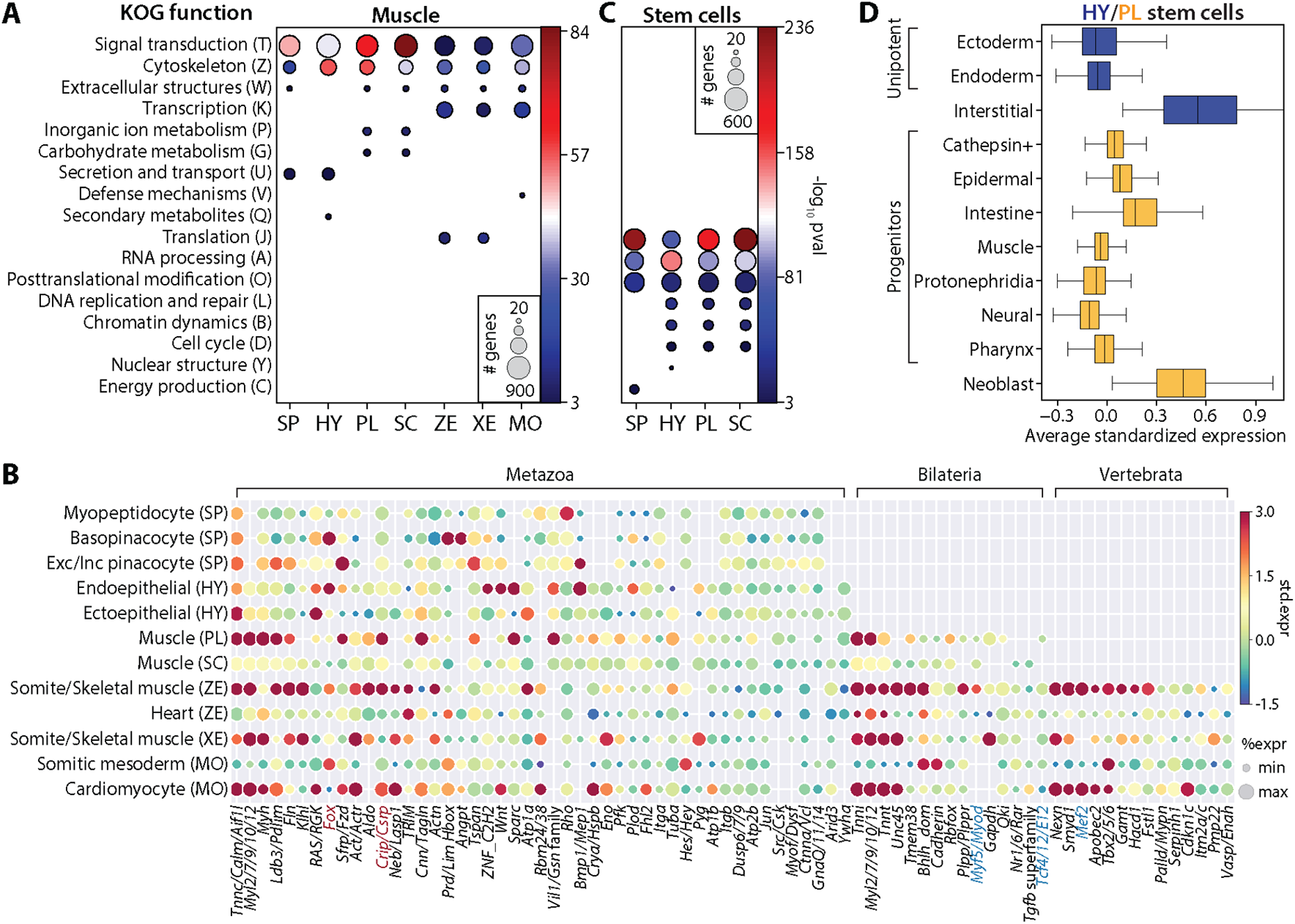
SAMap identifies muscle and stem cell transcriptional signatures conserved across species. (A) Enrichment of KOG functional annotations calculated for genes shared in contractile cell types. For each species, genes enriched in individual contractile cell types are combined. (B) Expression and enrichment of conserved muscle genes in contractile cell types. Color: mean standardized expression. Symbol size: the fraction of cells each gene is expressed in per cell type. Homologs are grouped based on overlapping eukaryotic Eggnog orthology groups. If multiple genes from a species are contained within an orthology group, the gene with highest standardized expression is shown. Genes in blue: core transcriptional program of bilaterian muscles; red: transcription factors conserved throughout Metazoa. (C) Enrichment of KOG functional annotations for genes shared by stem cell types. (D) Boxplot showing the median and interquartile ranges of the mean standardized expressions of genes in hydra and planarian stem cells/progenitors that are conserved across all invertebrate species in this study. Planarian progenitors: *piwi*+ cells that cluster with differentiated tissues in Fincher et al. (Fincher et al., 2018). Neoblasts: cluster 0 in Fincher et al. (Fincher et al., 2018) that does not express any tissue-specific markers.

We also identified several transcriptional regulators shared among contractile cells (**Figure 5B**). Previously known core regulators involved in myocyte specification (Brunet et al., 2016) were enriched only in bilaterian (e.g., *myod*, and *tcf4/E12*) or vertebrate contractile cells (e.g., *mef2*). In contrast, we found homologs of Muscle Lim Protein (*Csrp*) and Forkhead Box Group 1 (Larroux et al., 2008) enriched in contractile cells from all seven species. The Fox proteins included FoxC, which is known to regulate cardiac muscle identity in vertebrates (Brunet et al., 2016) and is contractile-specific in all species except schistosome and *Spongilla*. Notably, we also identified FoxG orthologs to be enriched in three of the four invertebrates (**Figure 5 – figure supplement 1**), suggesting that FoxG may play an underappreciated role in contractile cell specification outside vertebrates.

For the family of invertebrate stem cells, we identified 3,343 genes that are enriched in at least one cell type pair and observed significant enrichment (p-value < 10^−3^) of genes involved in translational regulation such as RNA processing, translation, and post-translational modification (**Figure 5C**). These genes form 979 orthology groups, 17% of which are enriched in all cell types of this family (**Supplementary Table 4**). Importantly, other stem cell populations in *Hydra* and planarian lineage-restricted neoblasts have significantly reduced expression of these genes (**Figure 5D**). These results suggest that SAMap identified a large, deeply conserved gene module specifically associated with multipotency.

## Discussion

Cell types evolve as their gene expression programs change either as integrated units or via evolutionary splitting that results in separate derived programs. While this notion of coupled cellular and molecular evolution has gained significant traction in the past years, systematically comparing cell type-specific gene expression programs across species has remained a challenging problem. Here, we map single-cell atlases between evolutionarily distant species in a manner that accounts for the complexity of gene evolution. SAMap aligns cell atlases in two mutually reinforcing directions, mapping both the genes and the cells, with each feeding back into the other. This method allows us to identify one-to-one cell type concordance between animals in the same phylum, whereas between phyla, we observe interconnected cell types forming distinct families. These findings support the notion that cell types evolve via hierarchical diversification (Arendt et al., 2019), resulting in cell type families composed of evolutionarily related cell types sharing a regulatory gene expression program that originated in their common ancestor. One-to-one cell type homologies should exist only if no further cell type diversification has occurred since the speciation.

In parallel, SAMap systematically identifies instances where paralogs exhibit greater expression similarity than orthologs across species. Paralog substitution likely occurs due to differential loss or retention of cell type-specific expression patterns of genes that were duplicated in the common ancestor (Studer & Robinson-Rechavi, 2009). Alternatively, paralog substitutions could arise due to compensating upregulation of paralogs following a loss-of-function mutation acquired by an ortholog (El-Brolosy et al., 2019). While the analysis presented here focuses on comparisons between two species, incorporating multiple species into a single analysis that also accounts for their phylogenetic relatedness could enable determining the specific order of paralog substitutions, associated cell type diversification events, and the mechanism by which they arose. However, this would require cell atlases that consistently sample key branching points along the tree of life. Nevertheless, identifying lineage-specific paralog substitution signatures should be accessible in extensively studied vertebrate single-cell atlases, as the vertebrate clade is where existing data and knowledge are most concentrated.

Besides applications in evolutionary biology, we anticipate SAMap can catalyze the annotation of new cell atlases from non-model organisms, which often represents a substantial bottleneck requiring extensive manual curation and prior knowledge. Its ability to use the existing atlases to inform the annotation of cell types in related species will keep improving as more datasets become available to better sample the diversity of cell types. Moreover, our approach allows leveraging existing and forthcoming single-cell gene expression data to predict functionally similar gene homologs, which can serve as guideposts for mechanistic molecular studies.

## Materials and Methods

### Data and Code Availability

- The source code for SAMap is publicly available at Github (https://github.com/atarashansky/SAMap), along with the code to perform the analysis and generate the figures.
- The datasets analyzed in this study are detailed in **Supplementary Table 1** with their accessions, and annotations provided.

### The SAMap Algorithm

The SAMap algorithm contains three major steps: preprocessing, mutual nearest neighborhood alignment, and gene-gene correlation initialization. The latter two are repeated for three iterations, by default, to balance alignment performance and computational runtime. SAMap runs up to one hour on an average desktop computer for 200,000 total cells.

#### 1. Preprocessing

##### 1.1. Generate gene homology graph via reciprocal BLAST

We first construct a gene-gene bipartite graph between two species by performing reciprocal BLAST of their respective transcriptomes using *tblastx*, or proteomes using *blastp. tblastn* and *blastx* are used for BLAST between proteome and transcriptome. When a pair of genes share multiple High Scoring Pairs (HSPs), which are local regions of matching sequences, we use the HSP with the highest bit score to measure homology. Only pairs with E-value < 10^−6^ are included in the graph.

Here we define similarity using BLAST, though SAMap is compatible with other protein homology detection methods (e.g. HMMER (Eddy, 2008)) or orthology inference tools (e.g. OrthoClust (Yan et al., 2014) and Eggnog (Huerta-Cepas et al., 2019)). While each of these methods has known strengths and limitations, BLAST is chosen for its broad usage, technical convenience, and compatibility with low-quality transcriptomes.

We encode the BLAST results into two triangular adjacency matrices, *A* and *B*, each containing bit scores in one BLAST direction. We combine *A* and *B* to form a gene-gene adjacency matrix *G*. After symmetrizing *G*, we remove edges that only appear in one direction: 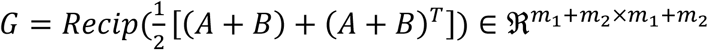, where *Recip* only keeps reciprocal edges, and *m*_1_ and *m*_2_are the number of genes of the two species, respectively. To filter out relatively weak homologies, we also remove edges where 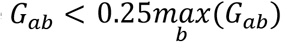. Edge weights are then normalized by the maximum edge weight for each gene and transformed by a hyperbolic tangent function to increase discriminatory power between low and high edge weights, 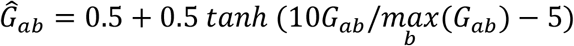.

##### 1.2. Construct manifolds for each cell atlas separately using the SAM algorithm

The scRNAseq datasets are normalized such that each cell has a total number of raw counts equal to the median size of single-cell libraries. Gene expressions are then log-normalized with the addition of a pseudocount of 1. Genes expressed (i.e.,*log*_2_(*D*+1) > 1)in greater than 96% of cells are filtered out. SAM is run using the following parameters: *preprocessing = ‘StandardScaler’, weight_PCs = False, k = 20*, and *npcs = 150*. A detailed description of parameters is provided previously (Tarashansky et al., 2019). SAM outputs *N*_1_and *N*_2_, which are directed adjacency matrices that encode *k*-nearest neighbor graphs for the two datasets, respectively.

SAM only includes the top 3,000 genes ranked by SAM weights and the first 150 principal components (PCs) in the default mode to reduce computational complexity. However, downstream mapping requires PC loadings for all genes. Thus, in the final iteration of SAM, we run PCA on all genes and take the top 300 PCs. This step generates a loading matrix for each species *i*, 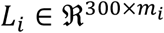.

#### 2. Mutual nearest neighborhood alignment

##### 2.1. Transform feature spaces between species

For the gene expression matrices 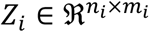, where *n* and *m* are the number of cells and genes respectively, we first zero the expression of genes that do not have an edge in *Ĝ* and standardize the expression matrices such that each gene has zero mean and unit variance, yielding 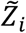. *Ĝ* represents a bipartite graph in the form of 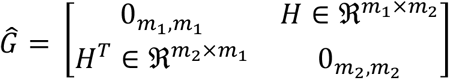, where 0_*m,m*_ is *m* × *m* zero matrix and *H* is the biadjacency matrix. Letting *H*_1_ = *H* and *H*_2_ = *H*^*T*^ encoding directed edges from species 1 to 2 and 2 to 1, respectively, we normalize the biadjacency matrix *H*_*i*_ such that each row sums to 1: 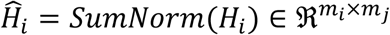, where the *SumNorm* function normalizes the rows to sum to 1. The feature spaces can be transformed between the two species via weighted averaging of gene expression, 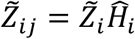

##### 2.2. Project single-cell gene expressions into a joint PC space

We project the expression data from two species into a joint PC space (Barkas et al., 2019), 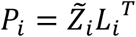 and 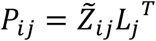. We then horizontally concatenate the principal components *P*_*i*_ and *P*_*ij*_ to form 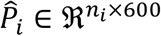.

##### 2.3. Calculate k-nearest cross-species neighbors for all cells

Using the joint PCs, 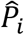, we identify for each cell the *k*-nearest neighbors in the other dataset using cosine similarity (*k* = 20 by default). Neighbors are identified using the *hnswlib* library, a fast approximate nearest-neighbor search algorithm (Malkov & Yashunin, 2020). This outputs two directed biadjacency matrices 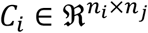 for (*i, j*) = (1,2) *or* (2,1) with edge weights equal to the cosine similarity between the PCs.

##### 2.4. Apply the graph-coarsening mapping kernel to identify cross-species mutual nearest neighborhoods

To increase the stringency and confidence of mapping, we only rely on cells that are *mutual* nearest cross-species neighbors, which are typically defined as two cells reciprocally connected to one another (Haghverdi et al., 2018). However, due to the noise in cell-cell correlations and stochasticity in the kNN algorithms, cross-species neighbors are often randomly assigned from a pool of cells that appear equally similar, decreasing the likelihood of mutual connectivity between individual cells even if they have similar expression profiles. To overcome this limitation, we integrate information from each cell’s local neighborhood to establish more robust mutual connectivity between cells across species. Two cells are thus defined as mutual nearest cross-species neighbors when their respective neighborhoods have mutual connectivity.

Specifically, the nearest neighbor graphs *N*_*i*_ calculated in step 1.2 are used to calculate the neighbors of cells *n*_*i*_ hops away along outgoing edges: 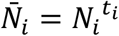, where 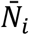 are adjacency matrices that contain the number of paths connecting two cells *n*_*i*_ hops away, for *i* = 1 *or* 2. *t*_*i*_ determines the length-scale over which we integrate incoming edges for species *i*. Its default value is 2 if the dataset size is less than 20,000 cells and 3 otherwise. However, cells within tight clusters may have spurious edges connecting to other parts of the manifold only a few hops away. To avoid integrating neighborhood information outside this local structure, we use the Leiden algorithm (Traag et al., 2019) to cluster the graph and identify a local neighborhood size for each cell (the resolution parameter is set to 3 by default). If cell *a* belongs to cluster *c*_*a*_, then its neighborhood size is *l*_*a*_ = |*c*_*a*_|. For each row *a* in 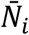 we only keep the *l*_*a*_ geodesically closest cells, letting the pruned graph be denoted as 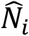

Edges outgoing from cell *a*_*i*_ in species *i* are encoded in the corresponding row in the adjacency matrix: 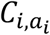. We compute the fraction of the outgoing edges from each cell that target the local neighborhood of a cell in the other species: 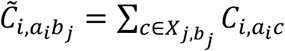 where 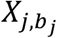 is the set of cells in the neighborhood of cell *b*_*j*_ in species *j* and 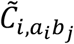 is the fraction of outgoing edges from cell *a*_*i*_ in species *j* targeting the neighborhood of cell *b*_*j*_ in species *j*.

To reduce the density of 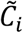 so as to satisfy computational memory constraints, we remove edges with weight less than 0.1. Finally, we apply the mutual nearest neighborhood criterion by taking the element-wise, geometric mean of the two directed bipartite graphs: 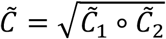 This operation ensures that only bidirectional edges are preserved, as small edge weights in either direction results in small geometric means.

##### 2.5. Assign the k-nearest cross-species neighborhoods for each cell and update edge weights in the gene homology graph

Given the mutual nearest neighborhoods 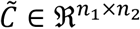, we select the *k* nearest neighborhoods for each cell in both directions to update the directed biadjacency matrices *C*_1_ and 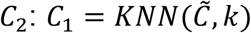 and 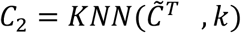, with *k* = 20 by default.

##### 2.6. Stitch the manifolds

We use *C*_1_ and *C*_2_ to combine the manifolds *N*_1_ and *N*_2_ into a unified graph. We first weight the edges in *N*_1_ and *N*_2_ to account for the number of shared cross-species neighbors by computing the one-mode projections of *C*_1_ and *C*_2_. In addition, for cells with strong cross-species alignment, we attenuate the weight of their within-species edges. For cells with little to no cross-species alignment, their within-species are kept the same to ensure that the local topological information around cells with no alignment is preserved.

Specifically, we use *N*_1_ and *N*_2_ to mask the edges in the one-mode projections, *Ñ*_1_ = *U(N*_*1*_)∘(*Norm*(*c*_1_*)Norm(c*_*2*_))and *Ñ*_*2*_ *= U(N*_*2*_)∘(*Norm*(*c*_2_*)Norm(c*_*1*_)),where *U*(*E*) sets all edge weights in graph *E* to 1 and *Norm* normalizes the outgoing edges from each cell to sum to 1. The minimum edge weight is set to be 0.3 to ensure that neighbors in the original manifolds with no shared cross-species neighbors still retain connectivity: *Ñ*_1,*ij*_ = *min(*0.3,*Ñ*_1,*ij*_) and *Ñ* _2_,_*ij*_ = *min(*0.3,*Ñ* _2,*ij*_)for all edges (*i, j*). We then scale the within-species edges from cell *i* by the total weight of its cross-species edges: 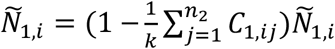 and 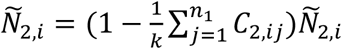. Finally, the within-and cross-species graphs are stitched together to form the combined nearest neighbor graph *N: N =*[*Ñ* _1_⊕ *C*_1_] ⊕ [*C*_2_ ⊕ *Ñ* _2_]. The overall alignment score between species 1 and 2 is defined as 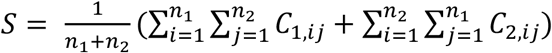.

#### 3. Gene-gene correlation initialization

##### 3.1. Update edge weights in the gene-gene bipartite graph with expression correlations

To compute correlations between gene pairs, we first transfer expressions from one species to the other: 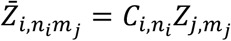, where 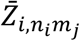 is the imputed expressions of gene *m*_*j*_ from species *j* for cell *n*_*i*_ in species *i*, and 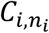 is row *n*_*i*_ of the biadjacency matrix encoding the cross-species neighbors of cell *n*_*i*_ in species *i*, all for (*i, j*) = (1,2) and (2,1). We similarly use the manifolds constructed by SAM to smooth the within-species gene expressions using kNN averaging: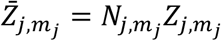, where *N*_*j*_ is the nearest-neighbor graph for species *j*. We then concatenate the within- and cross-species gene expressions such that the expression of gene *m*_*j*_from species *j* in both species is 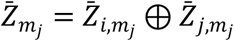 For all gene pairs in the unpruned homology graph generated in step 1.1., *Ĝ*, we compute their correlations, 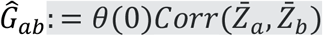, where *θ*(0) is a Heaviside step function centered at 0 to set negative correlations to zero. We then use the expression correlations to update the corresponding edge weights in *Ĝ*, which are again normalized through 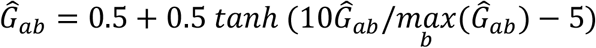.

### Annotation of cell atlases

To annotate the primary zebrafish and *Xenopus* cell types, the cell subtype annotations provided by the original publications (Briggs et al., 2018; Wagner et al., 2018) are coarsened using a combination of the manual matching and developmental hierarchies. For example, as “heart - mature”, “heart - hoxd9a”, “heart”, and “heart field” in zebrafish are all manually matched to “cardiac mesoderm” in *Xenopus*, we label these cells as “heart”. In cases where the matching is insufficient to coarsen the annotations, we use the provided developmental trees to name a group of terminal cell subtypes by their common ontogenic ancestor. The annotations provided by their respective studies were used to label the cells in the *Spongilla, Hydra*, and mouse atlases. To annotate the schistosome cells, we used known marker genes to annotate the main schistosome tissue types (Li et al., 2020). Annotations for all single cells in all datasets are provided in **Supplementary Table 1**.

### Visualization

The combined manifold *N* is embedded into 2D projections using UMAP implemented in the scanpy package (Wolf et al., 2018) by *scanpy*.*tl*.*umap* with the parameter *min_dist* = 0.1. The sankeyD3 package (https://rdrr.io/github/fbreitwieser/sankeyD3/man/sankeyD3-package.html) in R is used to generate the sankey plots. Edge thickness corresponds to the alignment score between mapped cell types. The alignment score between cell types *a* and *b* is defined as 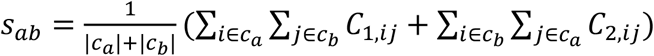 where *c*_*a*_ and *c*_*b*_ are the set of cells in cell types *a* and *b*, respectively. Cell type pairs with alignment score less than *z* are filtered out. By default, *z* is set to be 0.1. Cell types that did not cluster properly in their respective manifolds were omitted from the sankey plot. In the zebrafish-*Xenopus* comparison, we excluded heart, germline, and olfactory placode cells from both species because they did not cluster in the *Xenopus* atlas. Similarly, the iridoblast, epiphysis, *nanog*+, apoptotic-like, and forerunner cells were excluded because they did not cluster in the zebrafish atlas.

The network graphs in **Figure 4D** are generated using the *networkx* package (https://networkx.github.io) in python. To focus on densely connected cell type groups, we filter out cell type pairs with alignment score less than 0.05.

### Identification of gene pairs that drive cell type mappings

We define *g*_1_ and *g*_2_ to contain SAMap-linked genes from species 1 and 2, respectively. Note that a gene may appear multiple times as SAMap allows for one-to-many homology. Let 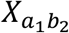 denote the set of all cells with cross species edges between cell types *a*_1_ and *b*_2_. We calculate the average standardized expression of all cells from species *i* that are in 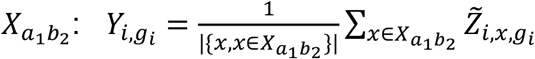, where 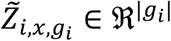 is the standardized expression of genes *g*_*i*_ in cell *x*. The correlation between 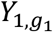 and 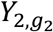 be written as 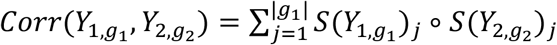 where *S*(*Z*) standardizes vector *Z* to have zero mean and unit variance. We use the summand to identify gene pairs that contribute most positively to the correlation. We assign each gene pair a score: 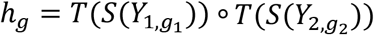, where *T*(*Z*) sets negative values in vector *Z* to zero in order to ignore lowly-expressed genes. To be inclusive, we begin with the top 1,000 gene pairs according to *h*_*g*_ and filter out gene pairs in which one or both of the genes are not differentially expressed in their respective cell types (p-value > 10^−2^), have less than 0.2 SAM weight, or are expressed in fewer than 5% of the cells in the cluster. The differential expression of each gene in each cell type is calculated using the Wilcoxon rank-sum test implemented in the *scanpy* function *scanpy*.*tl*.*rank_genes_groups*.

### Orthology group assignment

We used the Eggnog mapper (v5.0) (Huerta-Cepas et al., 2019) to assign each gene to an orthology group with default parameters. For the zebrafish-to-*Xenopus* mapping, genes are considered paralogs if they map to the same eukaryotic orthology group and orthologs if they map to the same vertebrate orthology group. For the pan-species analysis, we group genes from all species with overlapping orthology assignments. In **Figure 5B**, each column corresponds to one of these groups. As each group may contain multiple genes from each species, we present the expression of the gene with the highest enrichment score per species. All gene names and corresponding orthology groups are reported in **Supplementary Table 4**.

### Phylogenetic reconstruction of gene trees

We generate gene trees to validate the identity of genes involved in putative examples of paralog substitution and of *Fox* and *Csrp* transcriptional regulators that are identified as enriched in contractile cells. For this, we first gather protein sequences from potential homologs using the eggnog version 5.0 orthology database (Huerta-Cepas et al., 2019). For the *Fox* and *Csrp* phylogenies, we include all Fox clade I (Larroux et al., 2008) and Csrp/Crip homologs, respectively, from the seven species included in our study.

Alignment of protein sequences is performed with Clustal Omega version 1.2.4 using default settings as implemented on the EMBL EBI web services platform (Madeira et al., 2019). Maximum likelihood tree reconstruction is performed using IQ-TREE version 1.6.12 (Nguyen et al., 2015) with the ModelFinder Plus option (Kalyaanamoorthy et al., 2017). For the *Csrp* tree, we perform 1,000 nonparametric bootstrap replicates to assess node support. For *Fox*, we utilize the ultrafast bootstrap support option with 1,000 replicates. For each gene tree we choose the model that minimizes the Bayesian Information Criterion (BIC) score in ModelFinder. This results in selection of the following models: DCMut+R4 (*Csrp*) and VT+F+R5 (*Fox*). The final consensus trees are visualized and rendered using the ete3 v3.1.1 python toolkit (Huerta-Cepas et al., 2016) and the Interactive Tree of Life v4 (Letunic & Bork, 2019).

### KOG functional annotation and enrichment analysis

Using the eggnog mapper, KOG functional annotations are transferred to individual transcripts from their assigned orthology group. For enrichment analysis, all genes enriched in the set of cell type pairs of interest are lumped to form the target set for each species. For example, the target set for *Spongilla* archaeocytes used in **Figure 5C** is composed of all genes enriched between *Spongilla* archaeocytes and other invertebrate stem cells. Note that this set includes genes from other species that are linked by SAMap to the *Spongilla* archeocyte genes. We include genes from other species in the target set to account for differences in KOG functional annotation coverage between species. As such, the annotated transcripts from all 7 species are combined to form the background set. We used a hypergeometric statistical test (Eden et al., 2009) to measure the enrichment of the KOG terms in the target genes compared to the background genes.

### Mapping zebrafish and xenopus atlases using existing methods

For benchmarking, we used vertebrate orthologs as determined by Eggnog as input to Harmony (Korsunsky et al., 2019), Liger (Welch et al., 2019), Seurat (Stuart et al., 2019), Scanorama (Hie et al., 2019), BBKNN (Polański et al., 2019), which are all run with default parameters. One-to-one orthologs were selected from one-to-many and many-to-many orthologs by using the bipartite maximum weight matching algorithm implemented in *networkx*. When using the one-to-one orthologs as input for SAMap, we ran for only one iteration. The resulting integrated lower-dimensional coordinates (PCs for Seurat, Harmony, and Scanorama and non-negative matrix factorization coordinates for Liger) and stitched graph (BBKNN and SAMap) were all projected into 2D with UMAP (**Figure 2 – figure supplement 2A**). The integrated coordinates are used to generate a nearest neighbor graph using the correlation distance metric, which is then used to compute the alignment scores in **Figure 2 – figure supplement 2B**. The alignment scores for SAMap and BBKNN are directly computed from their combined graphs.

### In-situ hybridization in schistosomes

*S. mansoni* (strain: NMRI) juveniles are retrieved from infected female Swiss Webster mice (NR-21963) at ∼3 weeks post-infection by hepatic portal vein perfusion using 37°C DMEM supplemented with 5 % heat inactivated FBS. The infected mice are provided by the NIAID Schistosomiasis Resource Center for distribution through BEI Resources, NIH-NIAID Contract HHSN272201000005I. In adherence to the Animal Welfare Act and the Public Health Service Policy on Humane Care and Use of Laboratory Animals, all experiments with and care of mice are performed in accordance with protocols approved by the Institutional Animal Care and Use Committees (IACUC) of Stanford University (protocol approval number 30366). *In situ* hybridization experiments are performed as described previously (Tarashansky et al., 2019), using riboprobes synthesized from gene fragments cloned with the listed primers: collagen (Smp_170340): GGTGAAGAAGGCTGTTGTGG, ACGATCCCCTTTCACTCCTG; tropomyosin (Smp_031770): AAGCTGAAGTCGCCTCACTA, CATATGCCTCTTCACGCTGG; troponin (Smp_018250): CGTAAACCTGGTCAGAAGCG, ATCCTTTTCCTCCAGAGCGT; myosin regulatory light chain (Smp_132670): GAGACAGCGAGTAGTGGAGG, TGCCTTCTTTGATTGGAGCT; wnt (Smp_156540): TGTGGTGATGAAGATGGCAG, CCACGGCCACAACACATATT; frizzled (Smp_174350): CGAACAGGCGCATGACAATA, TGCTAGTCCTGTTGTCGTGT.

## Acknowledgments

We thank D. Wagner and C. Juliano for sharing the data and essential discussions. We also thank S. Granick, L. Luo, and J. Kebschull for their critical reading of the manuscript. AJT is a Bio-X Stanford Interdisciplinary Graduate Fellow. This work is supported by a Beckman Young Investigator Award and an NIH grant (1R35GM138061) to BW.

---

**Figure 2 – figure supplement 1:**
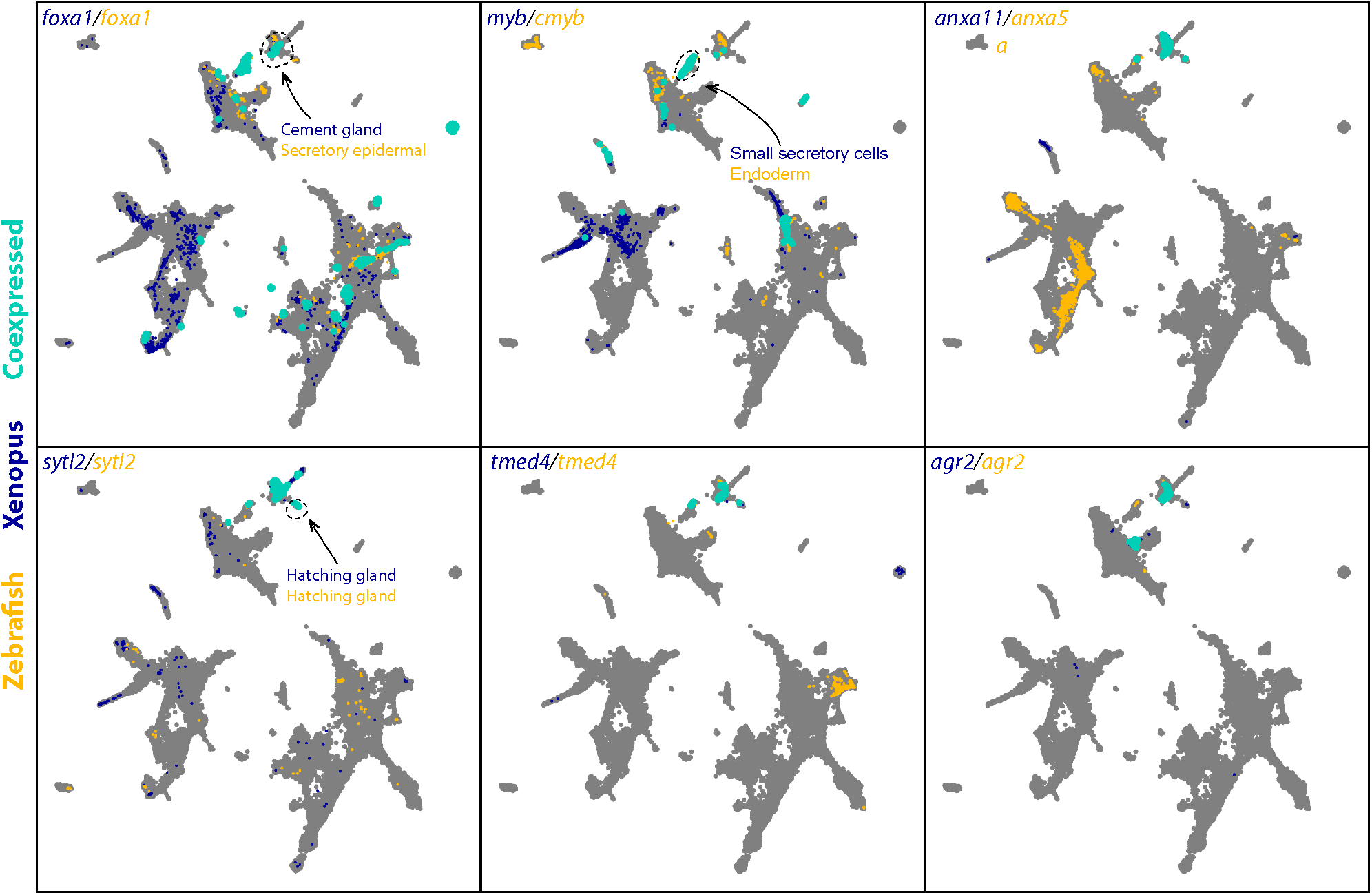
Expression of selected genes enriched in *D. rerio* and *X. tropicalis* secretory cell types. Expressions of orthologous gene pairs linked by SAMap are overlaid on the combined UMAP projection. Expressing cells are color-coded by species, with those connected across species colored cyan. Cells with no expression are shown in gray. The mapped secretory cell types are highlighted with circles.

**Figure 2 – figure supplement 2:**
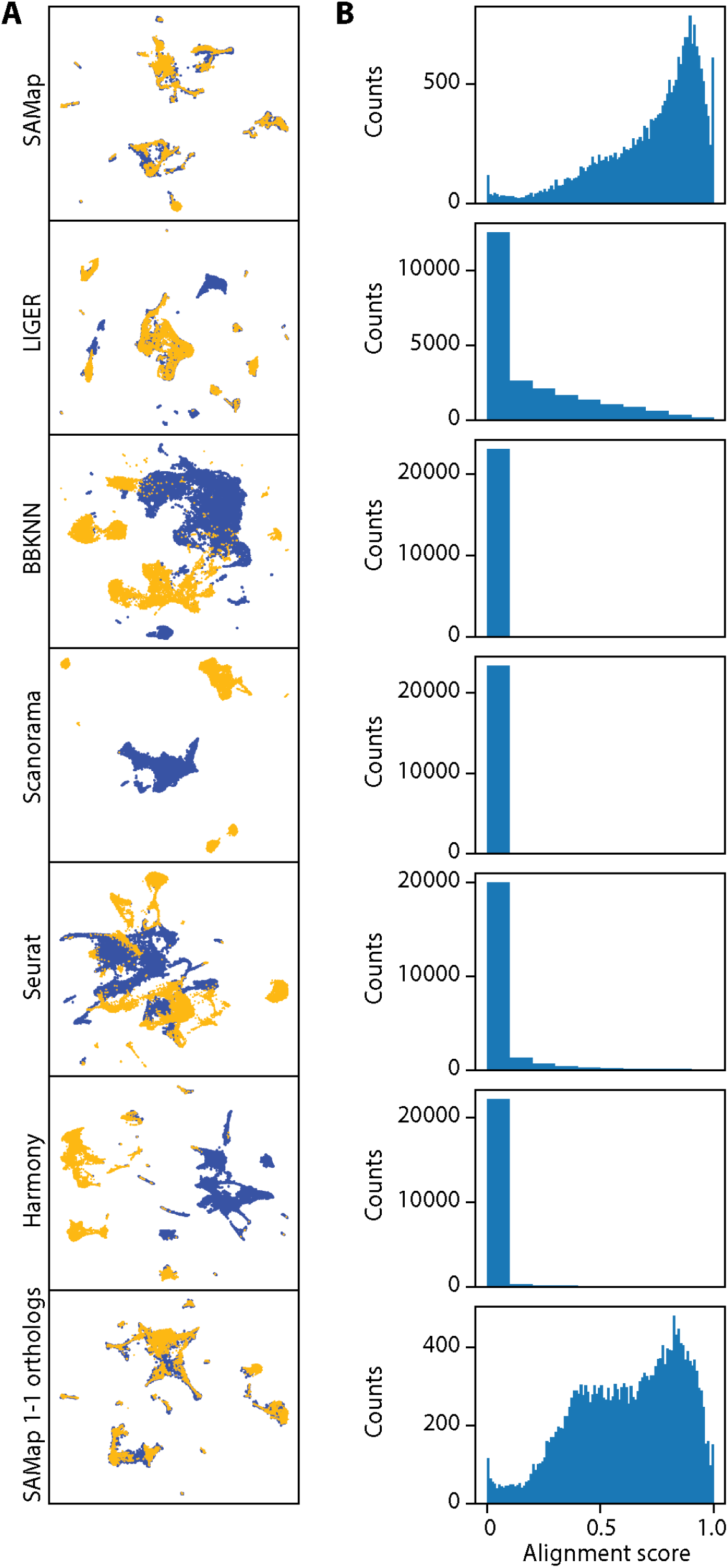
Existing methods failed to map *D. rerio* and *X. tropicalis* atlases. (A) UMAP projections of the integration results from SAMap using the full homology graph, compared to Liger, BBKNN, Scanorama, Seurat, Harmony, and SAMap using 1-1 orthologs. For fair comparisons, all methods were run on the *D. rerio* and *X. torpicalis* atlases subsampled to approximately 15,000 cells to satisfy computational constraints of Seurat and Liger. (B) Distribution of alignment scores between individual cells.

**Figure 2 – figure supplement 3:**
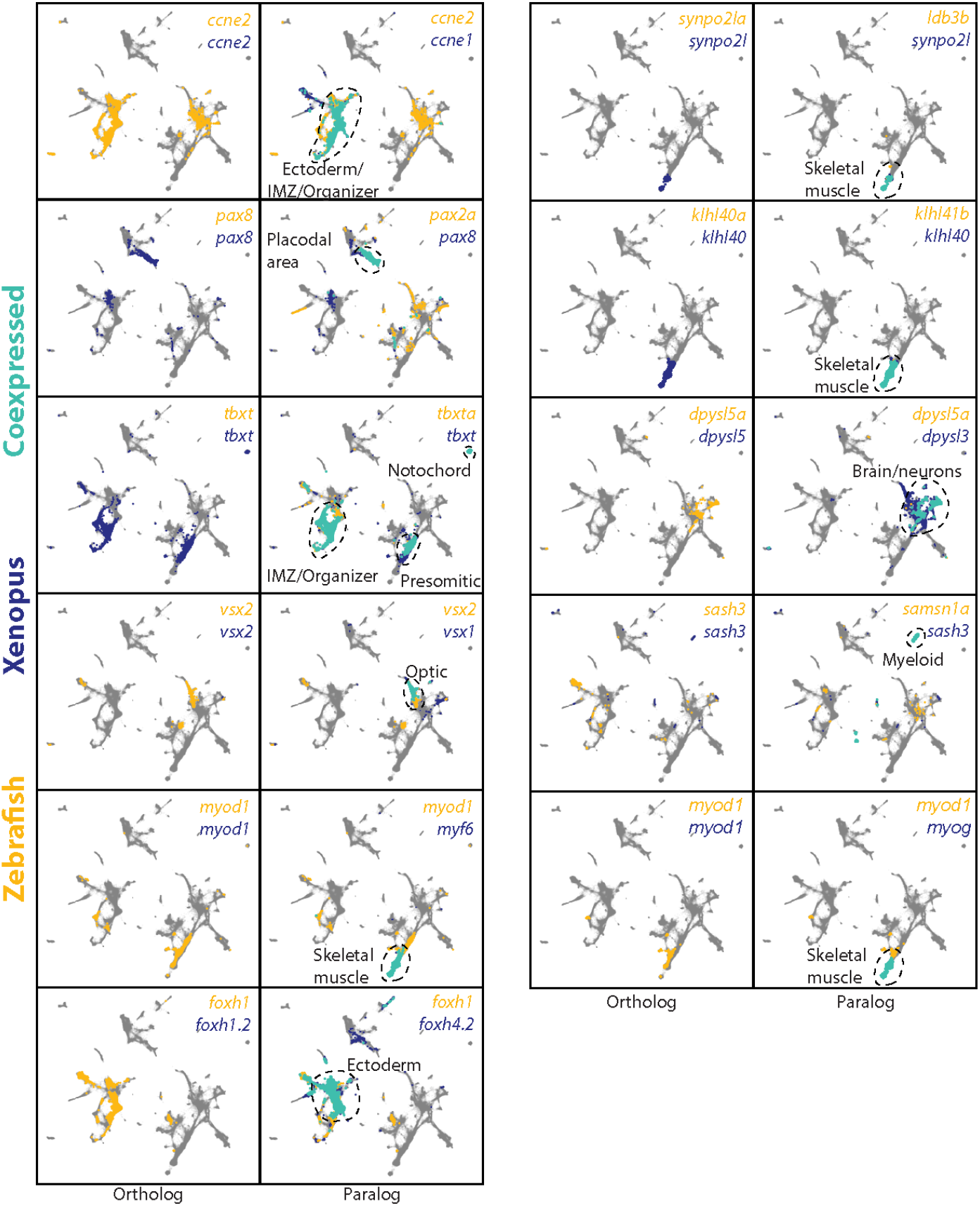
Representative examples of paralog substitution events in *D. rerio* and *X. tropicalis* atlases. Expressions of orthologous and paralogous gene pairs are overlaid on the combined UMAP projection. Expressing cells are color-coded by species, with those that are connected across species colored in cyan. Cells with no expression are shown in gray.

**Figure 3 – figure supplement 1:**
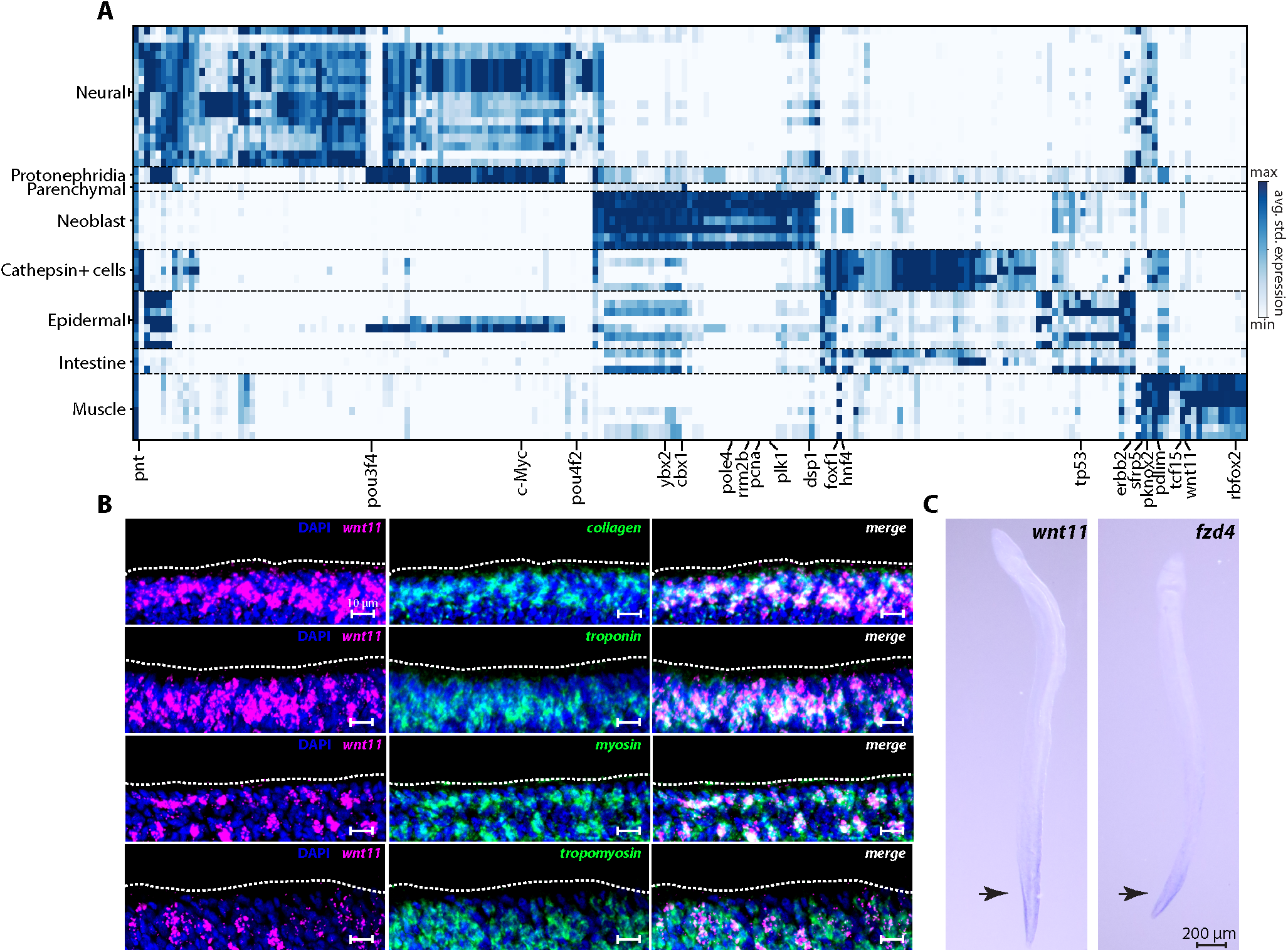
SAMap-linked gene pairs that are enriched in cell type pairs between *S. mediterranea* and *S. mansoni*. (A) Rows: linked cell types. Schistosome cell types correspond to leiden clusters. Columns: genes linked by SAMap with overlapping eukaryotic Eggnog orthology groups. We calculate the average standardized expression of each gene in an orthology group for its corresponding cell type in a particular pair and report the highest expression. A selected set of orthology groups corresponding to transcriptional regulators are labeled. (B) Fluorescence *in situ* hybridization shows the co-expression of *wnt* (Smp_156540) and a panel of muscle markers (*collagen, troponin, myosin* and *tropomyosin*) in *S. mansoni* juveniles. The body wall muscles are expected to be located close to the parasite surface (dashed outline). The images are maximum intensity projections constructed from ∼10 confocal slices with optimal axial spacing recommended by the Zen software collected on a Zeiss LSM 800 confocal microscope using a 40× (N.A. = 1.1, working distance = 0.62 mm) water-immersion objective (LD C-Apochromat Corr M27). (C) Whole mount *in situ* hybridization images showing that the expression of *wnt* and *frizzled* (Smp_174350) are concentrated in the parasite tail (arrows) with decreasing gradients extending anteriorly. In planarian muscles, Wnt genes provide the positional cues for setting up the body plan during regeneration (Scimone et al., 2017; Reddien, 2018). The presence of an anterior-posterior expression gradient of *wnt* and *frizzled* in muscles of schistosome juveniles suggests that they may have similar functional roles in patterning during development.

**Figure 3 – figure supplement 2:**
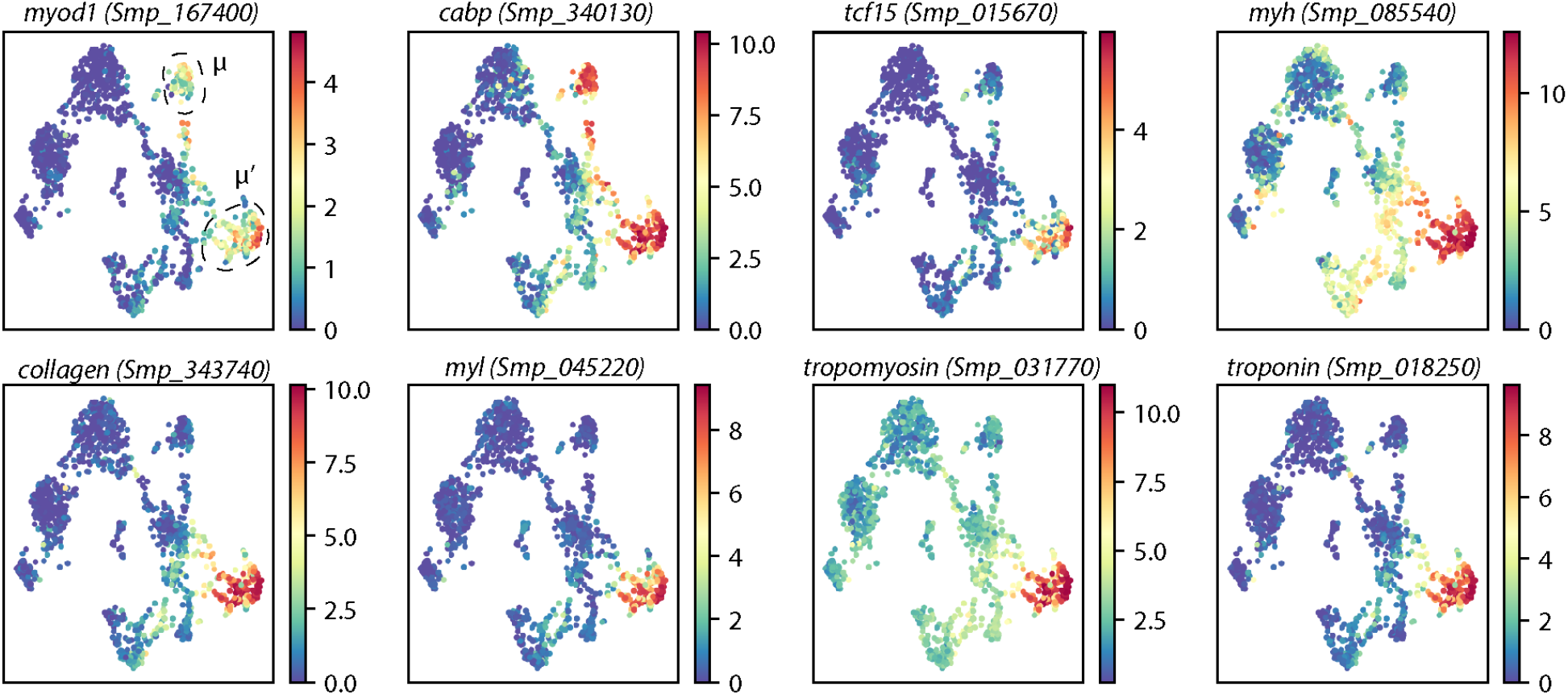
Schistosome neoblasts express canonical muscle markers in muscle progenitors. UMAP projections of schistosome neoblasts with gene expressions overlaid. μ and μ’ cells are circled. Colormap: expression in units of *log*_*2*_(*D+1*). For visualization, expression was smoothed via nearest-neighbor averaging using SAM. Note that *myod1* and *cabp* are expressed in both presumptive muscle progenitor populations, whereas all other markers are enriched in μ’ cells. All genes displayed are also expressed in fully differentiated muscle tissues.

**Figure 4 – figure supplement 1:**
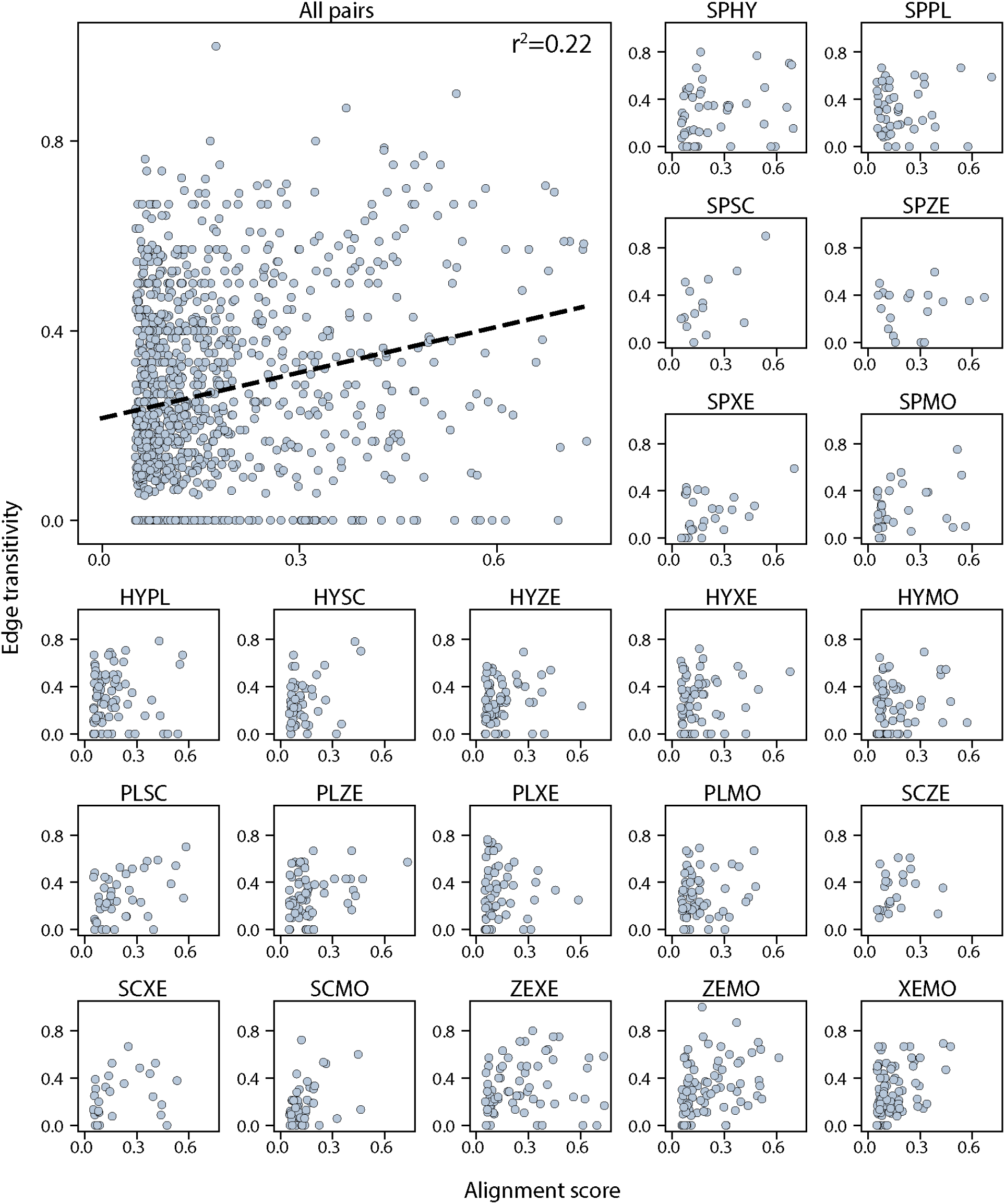
Alignment scores are mostly independent of edge transitivity. Top left: alignment scores and edge transitivity for all cell type pairs in the connectivity graph including the 7 species. Dotted line: the linear best fit, with the Pearson correlation coefficient reported at the top. Alignment scores and edge transitivity for individual species pairs are shown in the remaining subplots.

**Figure 4 – figure supplement 2:**
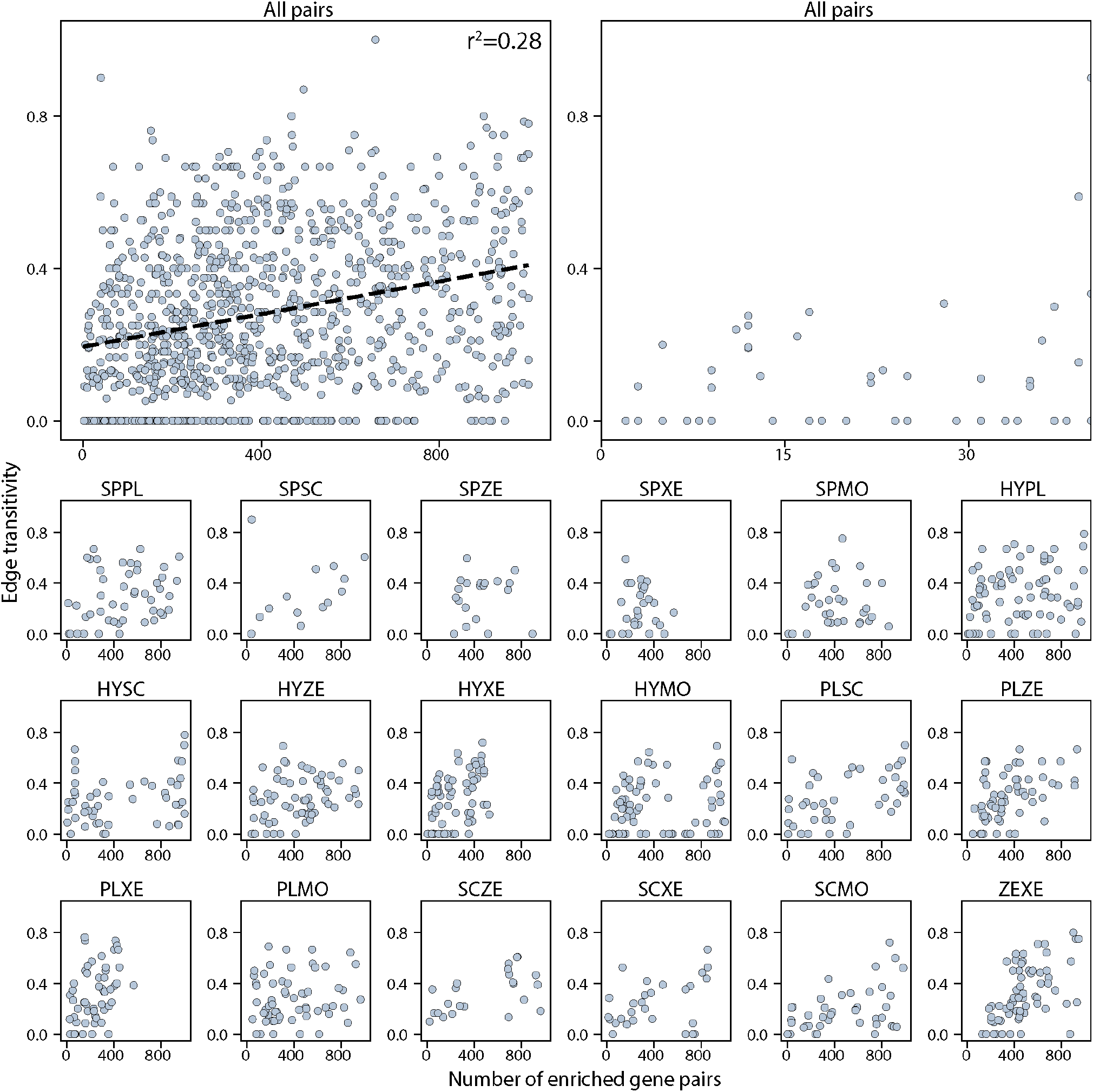
Number of enriched gene pairs are mostly independent of edge transitivity. Top left: The edge transitivity is plotted against the number of enriched gene pairs for all cell type pairs in the connectivity graph. Dotted line: the linear best fit, with the Pearson correlation coefficient reported at the top. Top right: magnified view of the mapped cell type pairs supported by small numbers of gene pairs (<40) to show those edges have low transitivity scores (<0.4). The sublots below show the number of enriched gene pairs and edge transitivity for individual species pairs.

**Figure 5 – figure supplement 1:**
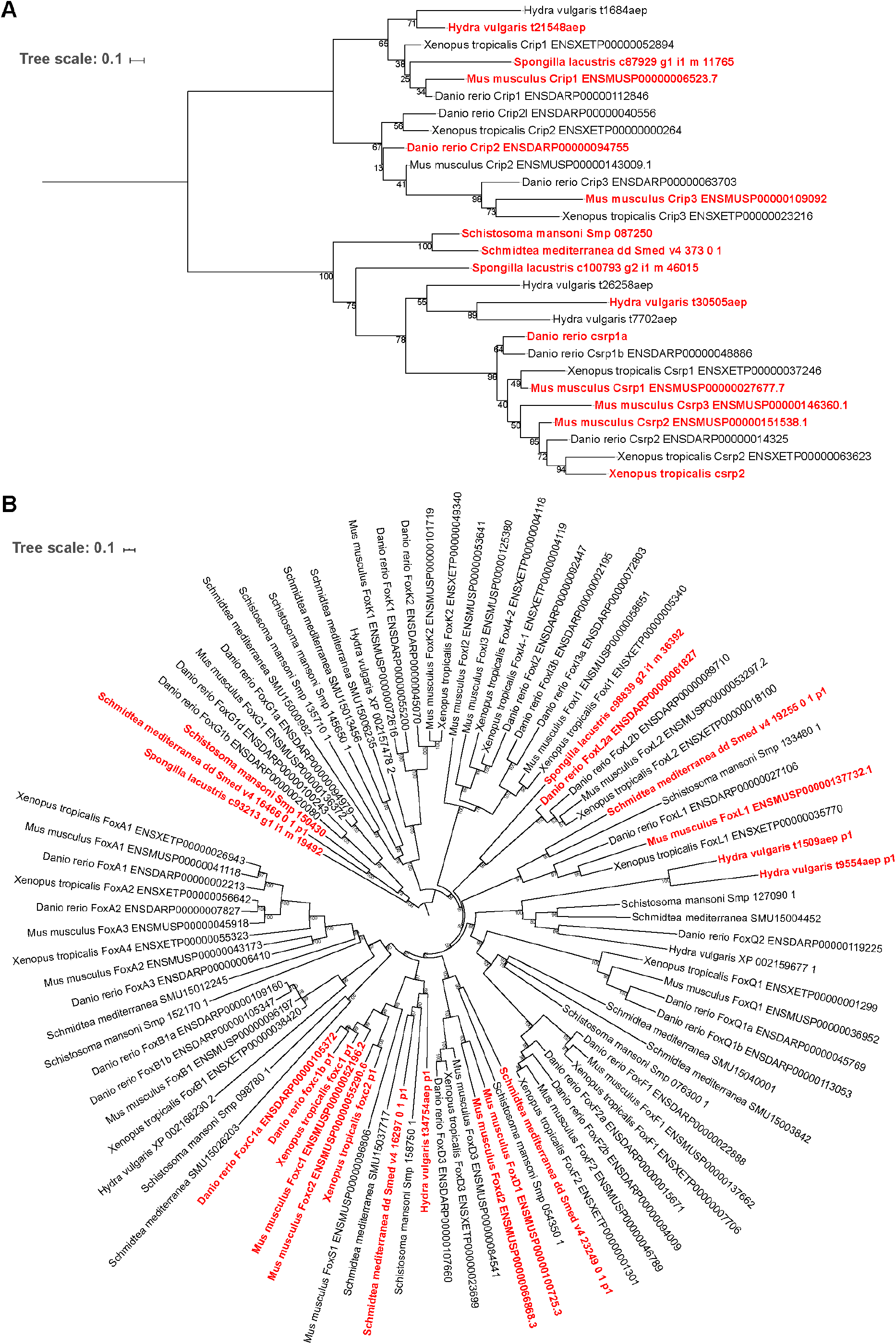
Phylogenetic reconstruction of animal contractile cell transcriptional regulators. Trees depict *Csrp/Crip* (A) and Fox group I (B) gene families. Genes labelled red are enriched in at least one contractile gene pair identified via SAMap. Support values indicate bootstrap support from 1,000 nonparametric (*Csrp*) or ultrafast (*Fox*) bootstrap replicates. Besides these two transcriptional regulators, contractile cells in all seven species were found to be also enriched for transcription factors from the C2H2 Zinc Finger, Lim Homeobox, and Paired Homeobox families, although in different cell types we found enrichment of a number of distinct orthologs. Whether this reflects an ancestral role for these transcription factor families in regulating contractility or their independent evolution will require additional taxonomic sampling and broader coverage of muscle cell diversity to resolve.

## Supplementary table captions

**Supplementary Table 1: Cell atlas metadata and cell annotations**. Metadata includes the number of cells, number of transcripts in the transcriptome, median number of transcripts detected per cell, the reference transcriptome used in this study, database through which the transcriptomes are provided, technology used for constructing the cell atlases, atlas data accessions, processing notes, and references. Leiden clusters and cell type annotations are reported for cells in each atlas. The Zebrafish and *Xenopus* tables include both the original cell type annotations and those used in this study. *D. rerio, X. tropicalis*, and mouse annotations include developmental stages.

**Supplementary Table 2: Cell type annotations for the zebrafish-*Xenopus* mapping**. Correspondence between the cell type annotations provided in the original study (Briggs et al., 2018; Wagner et al., 2018) and corresponding annotations used in this study is provided for both *D. rerio* and *X. tropicalis* atlases.

**Supplementary Table 3: Identified paralogs with greater expression similarity than orthologs in the zebrafish-*Xenopus* mapping**. Each row contains a pair of vertebrate-orthologous genes and a corresponding pair of eukaryotic paralogs with higher correlation in expression compared to the orthologs, the expression correlations for ortholog and paralog pairs, the difference between their correlations, and whether the paralogs are considered as a paralog substitution (defined as when the substituted ortholog is either absent or lowly-expressed with no cell-type specificity). Highlighted rows are shown in **Figure 2E** and **Figure 2 – figure supplement 3**.

**Supplementary Table 4: Genes enriched in contractile cell types and invertebrate stem cells highlighted in Figure 4D**. The IDs of the genes enriched in the contractile and invertebrate stem cell types are provided along with the IDs of the Eggnog orthology groups to which they belong. In cases where multiple genes from a species belonging to the same orthology group are enriched, the most differentially expressed gene is shown. The descriptions in the stem cell table are orthology annotations associated with the *Spongilla* genes provided in the original study (Musser et al., 2019).

**Supplementary Table 5: Cell types in the cell type families shown in Figure 4D**. For the schistosome cell types, we annotated two neural clusters, both of which express the neural marker *complexin* (Li et al., 2020). One of the clusters expresses the antigen *SmKK7*, so we label the clusters “Neural” and “Neural_KK7”, respectively. The “Muscle” population contains non-neoblast cells expressing *troponin*. The “Tegument_prog” and “Tegument” populations consist of cells expressing tegument progenitor and differentiated marker genes, respectively, as reported in a previous study (Wendt et al., 2018).

